# Cooperativity and folding kinetics in a multi-domain protein with interwoven chain topology

**DOI:** 10.1101/2022.02.02.478901

**Authors:** Zhenxing Liu, D. Thirumalai

**Affiliations:** Department of Physics, Beijing Normal University, Beijing 100875, China; Department of Chemistry, The University of Texas at Austin, Austin, TX 78712

## Abstract

Although a large percentage of eukaryotic proteomes consist of proteins with multiple domains, not much is known about their assembly mechanism, especially those with complicated native state architectures. Some have complex topology in which the structural elements along the sequence are interwoven in such a manner that the domains cannot be separated by cutting at any location along the sequence. We refer to such proteins as Multiply connected Multidomain Proteins (MMPs). The phoshotransferase enzyme Adenylate Kinase (ADK) with three domains (NMP, LID, and CORE), the subject of this study, is an example of MMP. We devised a coarse-grained model to simulate ADK folding initiated by changing either the temperature or guanidinium chloride (GdmCl) concentration. The simulations reproduce the experimentally measured melting temperatures that are associated with two equilibrium transitions, FRET efficiency as a function of GdmCl concentration, and the global folding times nearly quantitatively. Although the NMP domain orders independently, cooperative interactions between the LID and the CORE domains are required for complete assembly of the enzyme. The kinetic simulations show that on the collapse time scale, but less than the global folding time, multiple interconnected metastable states are populated, attesting to the folding heterogeneity. The network connectivity between distinct states shows that the CORE domain folds only after the NMP and LID domains are formed, reflecting the interwoven nature of the chain topology. We propose that the rules for MMP folding must also hold for the folding of RNA enzymes.

## Introduction

It is estimated that nearly seventy percent of eukaryotic proteins consist of multiple domains.^1^ They are involved in a wide array of functions, such as allosteric signaling (for example hemoglobin and the bacterial chaperonin GroEL), passive elasticity of muscle (titin), and cargo transport by motors (dynein). Despite the inherent difficulties in identifying domains in proteins, a perusal of their structures shows that there is a great deal of diversity in their architectures.^1^ For example, the giant titin protein is a heteropolymer made of thousands of *β*-sheet single-domain immunoglobulin (Ig) proteins that are connected by linkers. Pioneering single molecule pulling experiments^2^ on the polyprotein Ig_*n*_ established that it unfolds one domain at a time. It is likely that refolding, upon force quench, also proceeds by the formation of the native state, one domain at a time. Therefore, titin might assemble by preformed monomer units. We refer to polyproteins, such as Ig_*n*_, as Simply connected Multidomain Proteins (SMPs) because they can be partitioned into individual subunits by merely excising the linkers. Another example of SMP, Ankyrin Repeat, is shown in Fig.1A (Protein Data Bank ID: 3TWT). The amino acid residues, and the associated secondary structural elements (SSEs) in the SMPs are “one-dimensionally contiguous”.^3^ In contrast, in Multiply connected Multidomain Proteins (MMPs) the sequences are intertwined in such a manner that their structures cannot be dissected into independently folding subunits. Thus, topologically the domains cannot be cut in such a manner that they follow the sequence in a continuous linear manner, as is the case in SMPs. An example of MMP is T4 Lysozyme whose folding cooperativity was shown, using pulling experiments,^4^ to reflect the discontinuity in the connectivity of the SSEs in which a portion of the N-terminal sequence is part of the C-terminal domain. The connectivity of domains in terms of sequence is even more complicated in Adenylate Kinase (ADK), the protein of interest in this study, shown in Fig.1B (Protein Data Bank ID: 4AKE). According to the Wetaufer^3^ classification, the domains in ADK are discontinuous with respect to the sequence, and the connectivity of the SSEs. The rules linking the topology of the folded state of the MMPs are hard to anticipate based solely on the connectivity of sequence and SSEs. The problem is exacerbated because, with the exception of very few studies,^4–6^ there is a paucity of detailed experimental studies that have dissected the folding pathways of MMPs.

**Figure 1:**
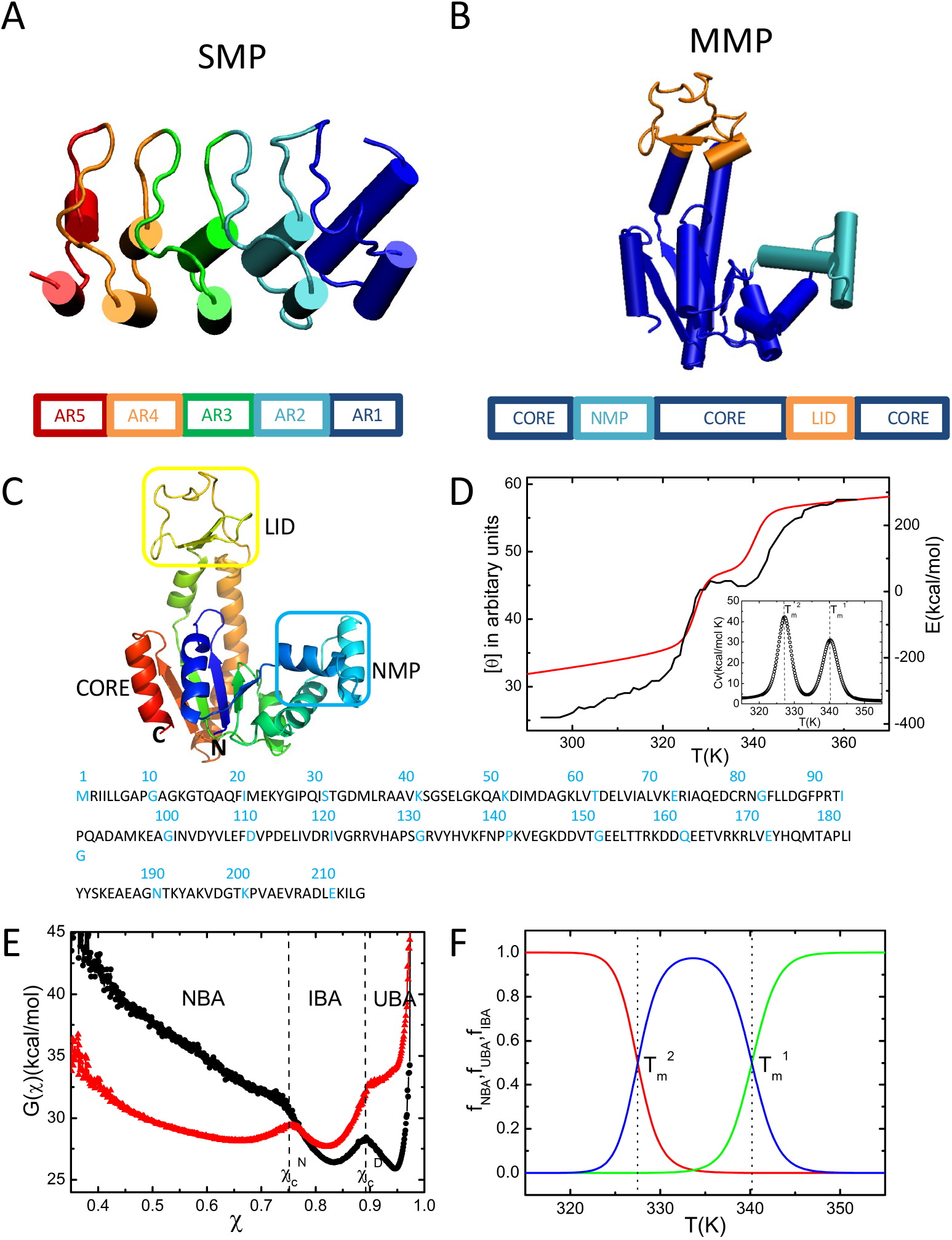
Structure, sequence and folding thermodynamics. (A) Cartoon representation of Ankyrin Repeat (PDB ID: 3TWT), an example for SMP. (B) Cartoon representation of ADK (PDB ID: 4AKE), an example for MMP. (C) Ribbon diagram representation of ADK. The **N** and **C** termini are indicated. The simulated sequence is shown below. (D) Temperature dependence of CD signal (black line), extracted from experiments^18^ and the simulated total energy (red line) as a function of temperature. The inset shows the calculated heat capacity as a function of temperature. (E) Free energy profiles at 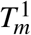 (black) and 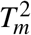 (red) as a function of the structural overlap function, *τ* (see Eq. (5)). The values 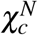 and 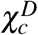 are used to separate the global equilibrium states, which are the Native Basin of Attraction (NBA), Intermediate Basin of Attraction (IBA), and Unfolded Basin of Attraction (UBA). (F) Temperature dependence of the fraction of molecules in the NBA (red), UBA (green), and IBA (blue).

In a series of most insightful experiments, Haas and coworkers reported the steps involved in the folding of *E. Coli*. ADK, triggered by varying guanidinium chloride (GdmCl) concentration.^6–8^ One of their key findings is that the collapse of ADK is fast but formation of secondary structural elements is slow.^7^ In a subsequent double kinetics experiments^6^ they further established that structure formation in the CORE domain (Fig.1B) is slow upon denaturant quench. More recently, Haran and coworkers used single-molecule fluorescence resonance energy transfer (sm-FRET) experiments^5^ to generate equilibrium trajectories at a fixed [*GdmCl*] concentration. Their results, which were analyzed Hidden Markov Model (HMM) analysis at different [*GdmCl*], suggest that folding occurs by the kinetic partitioning mechanism in which there are multiple metastable states in the folding landscape of ADK. The direct flux to the folded state from the unfolded ensemble (referred to as the partition factor elsewhere^9^) is only Φ ≈ 0.02,^10^ which implies that the majority of the molecules fold by first populating one of the (roughly six obtained from HMM analysis at 0.65M [*GdmCl*]) metastable states. Surprisingly, they found that connectivity between the states and the associated fluxes between them could be tuned by altering the denaturant concentration. Their experiments showed that not only is the folding landscape of ADK rugged but is also malleable to changes in the external conditions. Although qualitatively similar results have been found in the folding of PDZ3,^11^ a 110-residue protein with a much simpler topology than ADK, the intricate topology of the MMP renders the folding of the latter more complicated.

The structure of the 214-residue ADK consists of three domains: the NMP domain, the the LID domain and CORE domain (Fig.1C).^12^ The NMP domain spans residues 30-59 residues (indicated by the blue square in Fig.1C), the LID domain consists of residues 122-159 (indicated by the yellow square), and the rest of the residues (1-29, 60-121 and 160-214) belong to CORE domain. Sequence penetration across the native structure is vividly illustrated in ADK by noting that the N-terminal residues, 1-29, are part of the CORE domain comprising C-terminal residues. Contacts formed between N-terminal and C-terminal residues are labeled in the contact map of ADK in Fig.S1.

Here, we investigate thermal and denaturant-dependent folding of ADK using simulations based on the Self-Organized Polymer model with Side Chains (SOP-SC) and the Molecular Transfer Model (MTM),^13–15^ under conditions that closely mimic those used in the experiments.^5,6,10^ Coarse-grained model simulations, without side chains, were used to investigate folding cooperativity and multiple routes to the native state in ADK by thermal folding and unfolding.^16,17^ After demonstrating that our simulations quantitatively reproduce many experimental measurements, we show that cooperative interactions between the LID and CORE domains, with folding of the for-mer being slave to partial ordering of the latter, are required for ADK self-assembly. In contrast, the NMP domain folds independently at a higher temperature (or denaturant concentration) than the other two domains. The enhanced cooperative interactions between the LID and the CORE domains arises due to the discontinuous nature of the latter. The network of states linking the unfolded to the folded state, both at equilibrium and during refolding upon temperature quench, is multiply connected and shows that folding must occur by parallel pathways. The late stage of folding involves interaction between a reentrant helix in the CORE domain that forms contact with elements in N-terminal CORE domain. The methods used here are transferable for investigating the folding of other MMPs.

## Results

### Simulations predict thermal denaturation accurately

The Circular Dichroism (CD) spectrum shows that ADK undergoes two cooperative transitions, one at 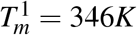, and the other at 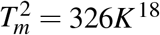 (Fig. 1D, black line). In order to assess if the simulations reproduce the observed thermal melting profile, we used Replica-Exchange Molecular Dynamics (REMD)^19–21^ and low friction Langevin dynamics^22^ in order to calculate the melting profiles. The temperature-dependent total energy *E*, which mirrors the CD signal, also shows two cooperative transitions (Fig.1D, red line). The corresponding melting temperatures, identified by the peaks in the heat capacity, *C*_*v*_, are at 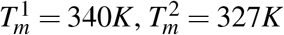 (see the inset of Fig.1D). The values of *T*_*m*_s obtained from our simulations and experiments are are in excellent agreement with each other. This is remarkable given that no parameter in the SOP-SC model was adjusted to obtain agreement with experiments.

### Three state folding

The temperature dependent profiles of *E* and *C*_*v*_ demonstrate that ADK folds globally in a three-state manner. The free energy profile, *G*(*χ*), as a function of the overlap function, *χ* given in Eq.(5), at 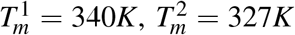 (Fig.1E), shows that three states, which represent the *NBA* (Native Basin of Attraction), *UBA* (Unfolded Basin of Attraction) and the *IBA* (Intermediate Basin of Attraction, i.e., *I*_*EQ*_). The conformations are grouped into three basins based on the *χ* values, shown by the black vertical dashed lines in Fig.1E. If 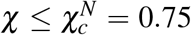, the conformations are classified as belonging to the NBA, conformations with 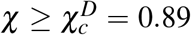 belong to the UBA, and the rest of the conformations represent IBA (*I*_*EQ*_). At 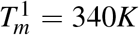, the NBA is unstable while *I*_*EQ*_ and the unfolded state have similar stabilities (Fig.1E, black line). At 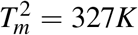, ADK transitions between the *NBA* and *IBA* while the *UBA* is unstable (Fig.1E, red line). Additional structural details of the *I*_*EQ*_ state are shown in Fig.S2.

In Fig.1F, we plot the fraction of molecules in the *NBA, f*_*NBA*_([0], *T*) (the first argument indicates the value of the denaturant concentration), in the *UBA, f*_*UBA*_([0], *T*), and in the *IBA, f*_*IBA*_([0], *T*). The temperature dependence of *f*_*UBA*_([0], *T*) (green curve in Fig.1F) and *f*_*NBA*_([0], *T*) (shown in red in Fig.1F) show that ADK unfolds and folds cooperatively at the two melting temperatures. At 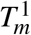, the value of *f*_*NBA*_([0], *T*) is negligible, reflecting the cooperative transition between the IBA and UBA. Using *f*_*IBA*_([0], *T*) = *f*_*UBA*_([0], *T*) = 0.5, we obtained 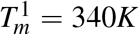, which coincides with the peak in the heat capacity (inset in Fig.1D). At low temperatures, the value of *f*_*UBA*_([0], *T*) is negligible, suggesting that ADK undergoes a cooperative transition between the NBA and the IBA. Using *f*_*IBA*_([0], *T*) = *f*_*NBA*_([0], *T*) = 0.5, we obtained 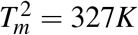, which also agrees with the peak in *C*_*v*_.

### Equilibrium folding of the domains

The average fraction of native contacts in each domain, *Q*^*NMP*^, *Q*^*LID*^, and *Q*^*CORE*^, as a function of temperature (Fig.2A), shows that the NMP and the LID domains fold in a two-state manner with the melting temperature, 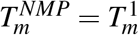, and, 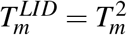. In contrast, the CORE domain folds in a three-state manner. The two melting temperatures, extracted from the temperature dependence of 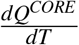, shows that the ordering of this domain reflects the two transition temperatures in the heat capacity and the total energy (Fig.1D). It follows that the incremental assembly of the CORE domain, across both the melting temperatures, is the reason that ADK globally folds in three stages.

**Figure 2:**
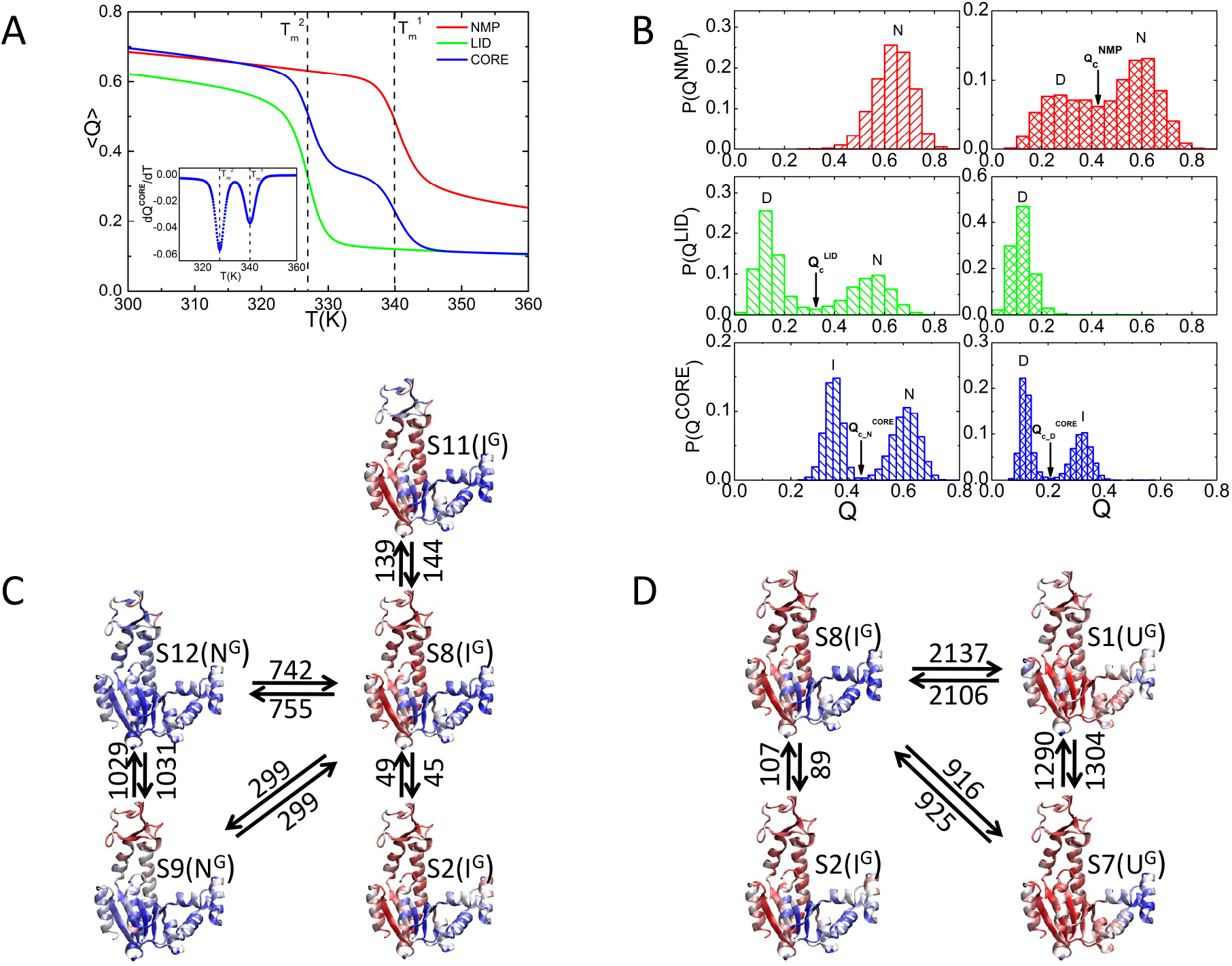
Temperature-dependent connectivity of metastable states. (A) Fraction of native contacts in each domain, *Q*^*NMP*^, *Q*^*LID*^, *Q*^*CORE*^ as a function of temperature. The inset shows the temperature dependence of 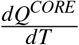. (B) The distributions of the fraction of native contacts within the three domains *P (Q*^*NMP*^, *P* (*Q*^*LID*^), and *P* (*Q*^*CORE*^_)_ at 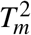 (left) and 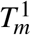 (right). (C) Network of thermodynamically connected substates at 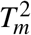. The numbers on the arrows are the transition times from one substate to another substate. (D) Same as (C) except it is calculated at 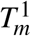.

A more nuanced picture of the folding thermodynamics emerges from the distributions of *Q*^*NMP*^, *Q*^*LID*^, *Q*^*CORE*^ at 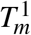and 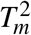 shown in Fig.2B. If 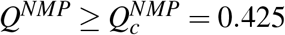, the NMP domain is predominantly folded 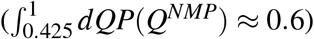, otherwise it is unfolded (compare the left and right panels in the upper panels in Fig.2B). The data in Fig.2A and Fig.2B show that the NMP domain forms before the LID and CORE domains become structured, as the temperature is decreased. Similarly, if 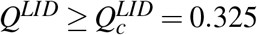, the LID substructure adopts native-like conformations (see the middle panels in Fig.2B). *P*(*Q*^*NMP*^) (*P*(*Q*^*LID*^)) is bimodal at 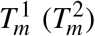 (Fig.2B), which is consistent with the interpretation that both the domains fold in an almost all-or-none manner, albeit at different melting temperatures. The lower melting temperature of the LID domain shows that it is thermodynamically less stable than the NMP domain, which accords well with single molecule pulling experiments.^23^ If 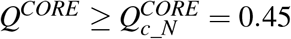, the CORE domain is in the native state. If the inequality 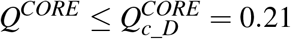 is satisfied, the CORE domain is in the unfolded state. In the intermediate state, we find that 0.21 *< Q*^*CORE*^ *<* 0.45 (see the bottom panels in Fig.2B). Two dimensional free energy profiles *G*(*Q*^*α*^, *Q*^*β*^) (*α* and *β* are the appropriate domain labels) in Fig.S3 illustrate the cooperativity between the LID and CORE domains at the two melting temperatures.

### Network of connected substates

The NMP and LID domains exhibit two state-like transitions as *T* is varied whereas the CORE domain ordering is best described using three states labeled as U, I, and N (Fig.2A). Thus, from a thermodynamic perspective we could describe the formation of ADK using 2 ×2 ×3 = 12 substates. They are S1(UUU), S2(UUI), S3(UUN), S4(UNU), S5(UNI), S6(UNN), S7(NUU), S8(NUI), S9(NUN), S10(NNU), S11(NNI), S12(NNN). The first letter in the brackets represents the state of the NMP domain, the second letter represents the state of the LID domain and the third letter stands for the state of the CORE domain.

We first determined the percentages of the substates in each global state obtained in the simulations by generating 28 folding trajectories. The conformations that are sampled were grouped into the 12 substates (S1-S12) and the 3 global states (*U*^*G*^, *I*^*G*^, and *N*^*G*^). The percentages are determined from the number of each substate in each global state. Out of the total 12 substates, only 7 substates are significantly populated. We find that the global native state is a superposition of the substate S9 (7.3%) and S12 (92.7%), which we write as *N*^*G*^ = 7.3% · *S*9 + 92.7% · *S*12. Similarly, the globally unfolded state is decomposed as *U*^*G*^ = 92.8% · *S*1 + 7.2% · *S*7. For the global intermediate state, we find *I*^*G*^ = 1.1% · *S*2 + 97.8% · *S*8 + 1.1% · *S*11.

We then performed a flux analysis among these substates at the two melting temperatures (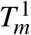 and 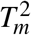) to assess the complexity of the network connectivity in thermodynamic folding landscape. At the lower melting temperature, 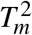, the equilibrium flux predominantly flows through the substates S9, S12 and S2, S8, S11 (Fig.2C). Considering their global structural features, the network shows that ADK transitions primarily between *N*^*G*^ and *I*^*G*^. The numbers on the arrows are the transition times from one substate to another substate. At the higher melting temperature 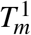, the network connectivity involves predominantly the substates S2, S8 and S1, S7(Fig.2D). By mapping to the global structural features, we find that ADK transitions back and forth between *I*^*G*^ and *U*^*G*^ at the higher melting temperature.

### Chemical Denaturation

In order to compare with experiments directly, we first used the Molecular Transfer Model (MTM)^13^ to simulate the effects of GdmCl on the equilibrium properties. Following our previous studies,^13–15^ we chose a simulation temperature, *T*_*s*_, at which the calculated free energy difference between the native state (*N*^*G*^) and the unfolded state (*U*^*G*^), Δ*G*_*NU*_(*T*_*s*_) 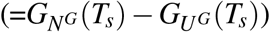 and the measured free energy Δ*G*_*NU*_ (*T*_*E*_) at *T*_*E*_ (=293K) coincide. The use of Δ*G*_*NU*_ (*T*_*s*_) = Δ*G*_*NU*_ (*T*_*E*_) (in water) to fix *T*_*s*_ is equivalent to choosing the overall reference free energy scale in the simulations. For ADK, Δ*G*_*NU*_ (*T*_*E*_ = 293*K*) = −9.8*kcal/mol* at [*C*] = 0,^18^ which results in *T*_*s*_ = 322*K*. Except for the choice of *T*_*s*_, no other parameter is adjusted to obtain agreement with experiments for any property.

With *T*_*s*_ = 322*K* fixed, we first computed the FRET efficiency as a function of [GdmCl] for ADK (Fig.3A). The FRET efficiency of a protein conformation was calculated using 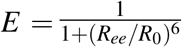, where *R*_*ee*_ is the end-to-end distance and *R*_0_ = 49 Å. In the experiments, residues 73, 203 were labelled.^24^ The agreement between the computed (thick black line) and the measured (black dots)^5^ FRET efficiencies is excellent. The derivative of the computed FRET efficiency with respect to [GdmCl] (inset of Fig.3A) also shows signs of the two thermodynamic transitions as the denaturant concentration is increased, which accords well with our thermal unfolding calculations. The midpoint concentration for the major transition is 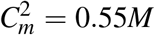, which agrees well with the measured result.^24^ The predicted midpoint concentration for the second transition is at 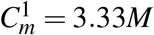, which has not been observed in experiments. The values for the FRET efficiency for the structures in the UBA are roughly constant as [*GdmCl*] changes (green line in Fig.3A) whereas the values for the FRET efficiency for the structures in the NBA decreases substantially as [*GdmCl*] increases (red line in Fig.3A).

**Figure 3:**
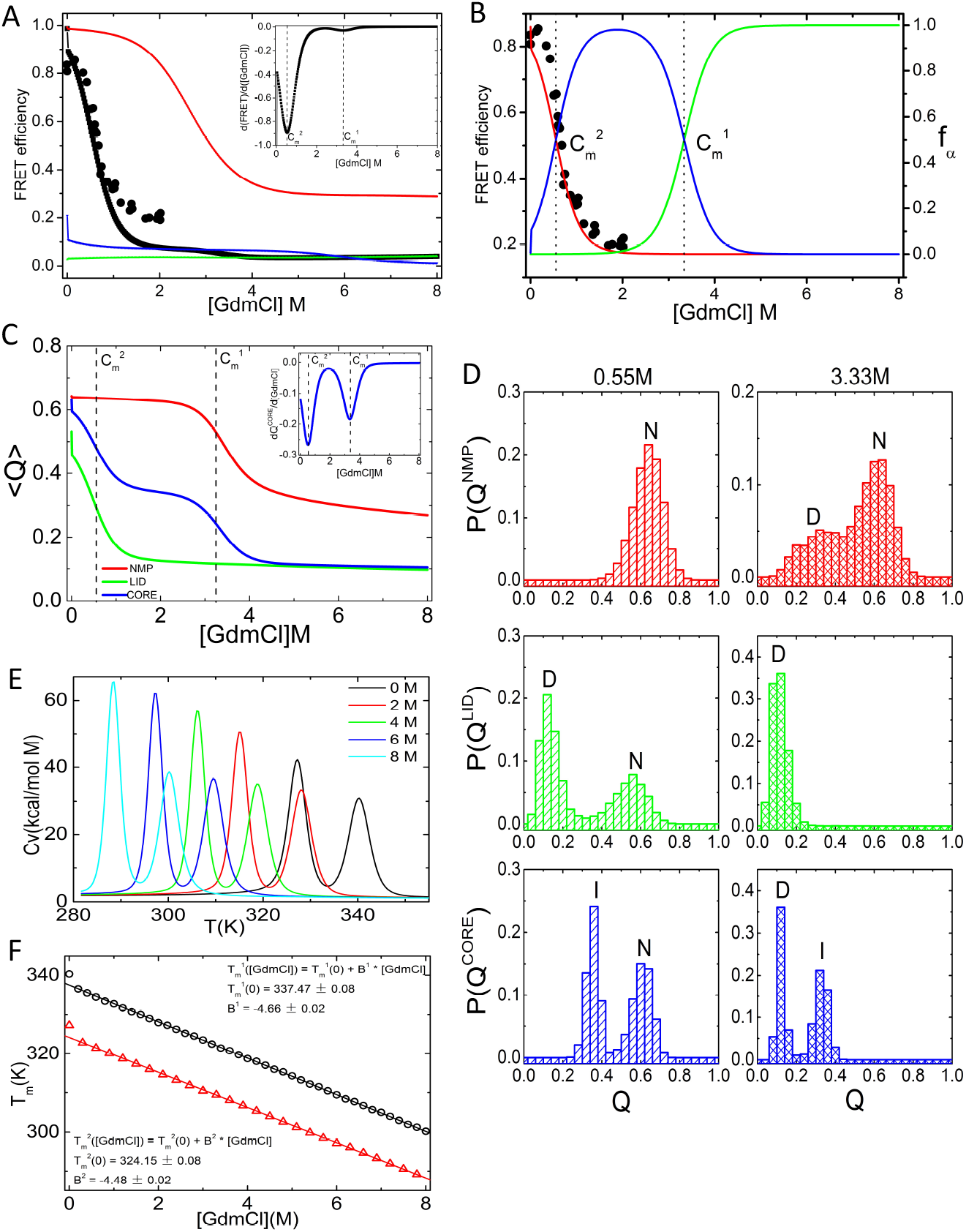
Effect of GdmCl on ADK folding. (A) Comparison of the calculated (thick black line) and experimental measurements^5^ (black dots) of the FRET efficiencies as a function of [*GdmCl*], the GdmCl concentration. The inset shows the derivative of the calculated FRET efficiency, which clearly indicates there are two distinct transitions. Decomposition of the FRET efficiency for the structures in the NBA (red), UBA (green), and IBA (blue). (B) Comparison of experimental measurements of the FRET efficiencies (black dots)^5^ and the calculated fraction of ADK molecules in the NBA (red), UBA (green), and IBA (blue) as a function of [*GdmCl*]. The comparison shows that below ≈ 0.8M the experimental FRET efficiency coincides with the calculated values for ADK molecules that are predominantly in the NBA, which is consistent with the plot in (A). In the range 0.8 ≤ [*GdmCl*] ≤ 2M molecules both the NBA and IBA contribute to the FRET efficiency. (C) Fraction of native contacts in each domain, *Q*^*NMP*^, *Q*^*LID*^, *Q*^*CORE*^ as a function of [GdmCl]. The inset shows the denaturant dependence of 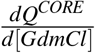. (D) The distributions of the fraction of native contacts within the three domains *P*(*Q*^*NMP*^), *P*(*Q*^*LID*^), and *P*(*Q*^*CORE*^) at 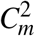 (left) and 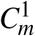 (right). (E) Heat capacity versus temperature for different values of [*GdmCl*]. (F) The [*GdmCl*] dependence of the melting temperatures. The fit to the lines are explicitly displayed. The units of *B*^1^ and *B*^2^ are K.M^−1^. The black (red) line is for 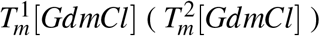.

The dependence of the simulated *f*_*NBA*_([*GdmCl*], *T*_*s*_) on [*GdmCl*] is also in excellent agreement with the measured FRET efficiency^5^ (Fig.3B). As in the case of thermal denaturation, the transition at low [*GdmCl*] takes place between the NBA and the IBA. The corresponding midpoint concentration 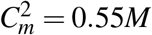, determined using *f*_*NBA*_([*GdmCl*], *T*_*s*_) = *f*_*IBA*_([*GdmCl*], *T*_*s*_) = 0.5 (red and blue lines in Fig.3B), is close to the experimental value.^24^ The transition at high [*GdmCl*] occurs as the intermediate state is destabilized, thus populating the unfolded state. Using *f*_*UBA*_([*GdmCl*], *T*_*s*_) = *f*_*IBA*_([*GdmCl*], *T*_*s*_) = 0.5 (green and blue lines in Fig.3B), the associated midpoint concentration is 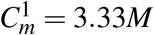.

The variations in the average values of the fraction of native contacts in the various domains (*Q*^*NMP*^, *Q*^*LID*^, and *Q*^*CORE*^), shown in Fig.3C, as a function of [GdmCl], are very similar to the results in Fig.2A. The distributions of *Q*^*NMP*^, *Q*^*LID*^, and *Q*^*CORE*^ at the two midpoint concentrations (Fig.3D) are also qualitatively similar to the ones calculated at the two melting temperatures (Fig.2B). However, there is a subtle difference. In the presence of the denaturant, the range of conformations that are accessed is broader. For example, the probability of sampling the ordered state of LID domain (⟨*Q*^*LID*^ ⟩ > 0.6) is non-negligible at 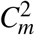 whereas it is much smaller at 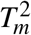 (compare the middle panels in Fig.2B and Fig.3D). This subtle difference could result in the differences in the stability of the folded ADK in the presence of GdmCl and folding induced by lowering the temperature.

The heat capacity curves at various values of [GdmCl] show that the peaks corresponding to 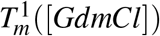 and 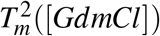 decrease as [*GdmCl*] increases (Fig.3E). The decrease in 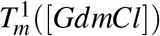 and 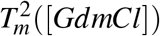 is both linear (Fig. 3F). The variation in 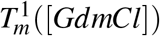 is well fit using 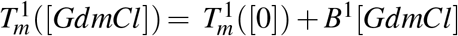, where *B*^1^ ≈ −4.7*K/M*. The variation in 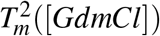 can be fit similarly using 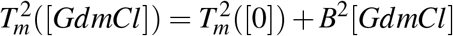, where *B*^2^ ≈ −4.5*K/M*.

### Collapse kinetics and folding kinetics

To analyze the collapse and folding kinetics, we generated 100 folding trajectories (see SI for details) using Brownian dynamics simulations at *T* = 293*K* at [*GdmCl*] = 0*M*.^25^ We calculated the time-dependent changes in the radius of gyration (⟨*R*_*g*_(*t*) ⟩ by averaging over the ensemble of trajectories). The decay of ⟨*R*_*g*_(*t*) ⟩, which is a measure of the extent of collapse, is fit using a single exponential function (Fig.4B), yielding collapse rate *k*_*c*_ = 391*s*^−1^, which as we discuss below is larger than the folding *k* _*f*_. Thus, global compaction occurs before folding, as observed in the experiment.^26^ In particular, the distances *d*(28 − 71)(*t*) and *d*(122 − 159) approach the native values extremely rapidly (Fig.4C), which likely corresponds to the dead time of the experiments.^26^ Fig.S5A and Fig.S5C in the SI show that the time for the probabilities of these two distances, *P*(*d*(28 − 71))(*t*) and *P*(*d*(122 − 159))(*t*), to exceed about 0.5 is ≈ 2ms, which is on the order of the collapse time 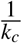. Thus, global compaction occurs rapidly upon making the conditions favorable for folding, as observed in the experiment.^26^

**Figure 4:**
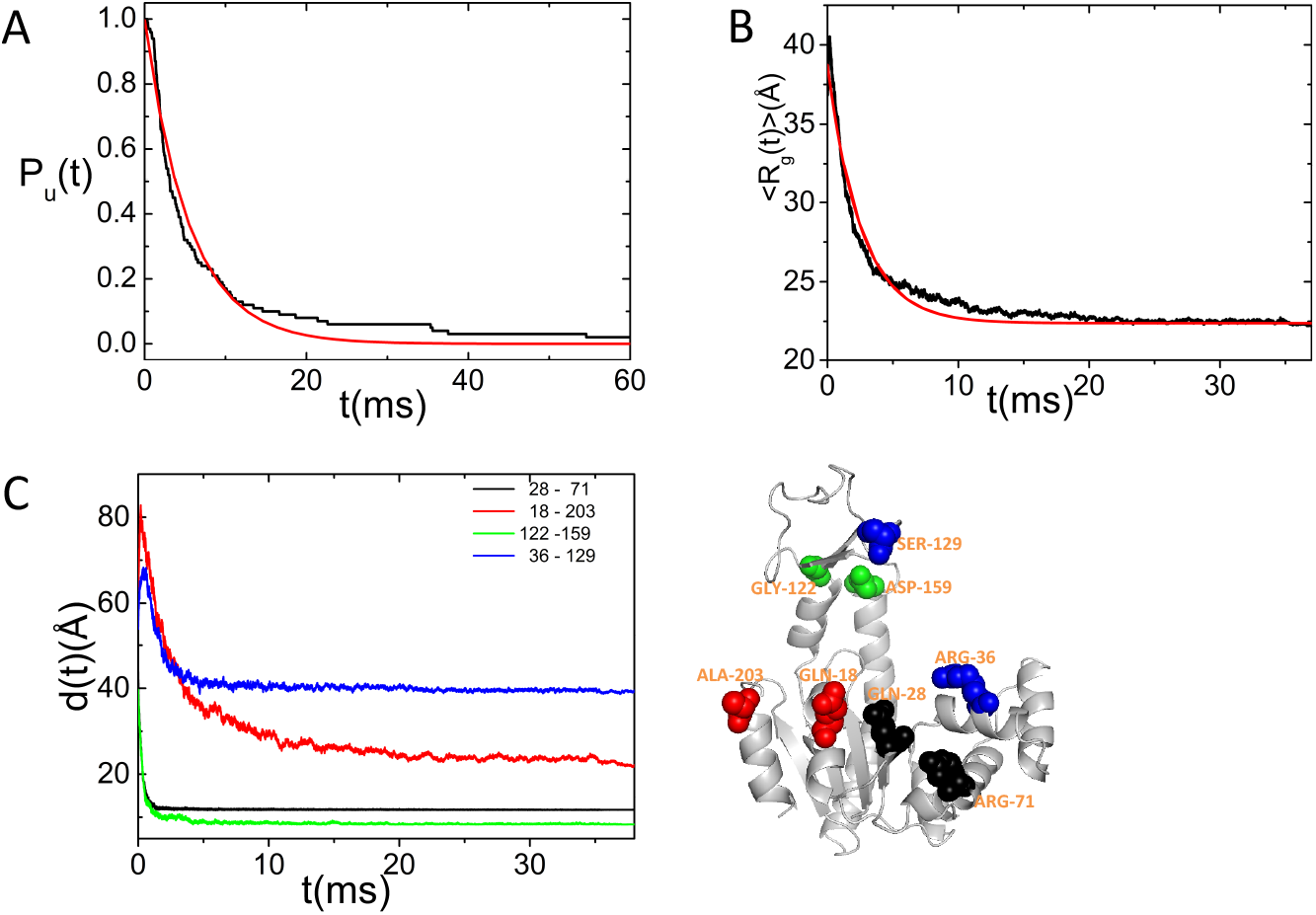
Folding and collapse kinetics. (A) Fraction of unfolded ADK molecules as a function of time (black) calculated from the distribution of first passage times. The red line is an exponential fit 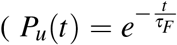 with *τ* = 5.5*ms*) to the data. (B) The kinetics of collapse monitored by the average *< R*_*g*_(*t*) *>* as a function of *t* (black). The fit to the data, given by a single exponential function (red line) yields an average collapse time, *τ*_*c*_ = 2.56*ms*. (C) The time dependent changes in the distances between residues 28 and 71 (black line), 18 and 203 (red line), 122 and 159 (green line), 36 and 129 (blue line). In the folded state, the distances between these four pairs of residues are 11.3Å, 13.1Å, 7.6Å, and 26.5Å, respectively.

Before estimating the folding time from simulations, we first calculated the folding rate theoretically using 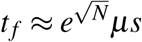,^27,28^ which gives fairly accurate estimates for the folding times, spanning nearly ten orders of magnitude, for proteins of varying length.^29^ For ADK, *N* = 214, we find that 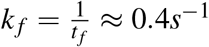, which is in good agreement with experiments.^7^ From the distribution of first passage times *P*_*f p*_(*s*), the fraction of unfolded molecules at time *t* is calculated using 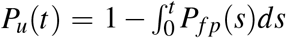. An exponential fit, 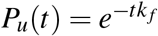, yielded the folding rate at [*GdmCl*] = 0*M, k* _*f*_ = 182*s*^−1^(Fig.4A). The calculated value is larger than the experimental value of ≈ (0.5 − 0.7)*s*^−1^ obtained by quenching the denaturant concentration from a high value to [GdmCl]=0.3M. The experimental [GdmCl] is fairly close to 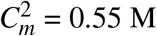. Therefore, one has to account for the stability change so that the simulation results could be compared to experiments directly. If the corrections due to stability change at 0.3M is made (Fig.S4), using the data in Fig.3B, we predict that *k* _*f*_ at [*GdmCl*] = 0.3*M* is ≈ 0.3*s*^−1^(Details in SI), which agrees well with measurements.^6,7^

### Heterogeneity in the self-assembly of ADK

The generated folding trajectories could be used to extract quantitatively the extent of folding heterogeneity. In particular, ensemble FRET experiments provide data on the time-dependent changes in FRET efficiencies by varying the positions of the FRET probes. We calculated the time dependence of the distances between four pairs of residues: (i) the distance between 28 and 71, which are at the ends of a 44-residue segment including the NMP domain. (ii) the distance between residues 18 and 203, which could be a reporter of the global folding, (iii) the distance between residues 122 and 159, which are the ends of LID domain, and (iv) the distance between residues 36 and 129, which reflects the closeness of the NMP and LID domains (Fig.4C). The time-dependent changes in these distances are well fit using single exponential functions, from which we obtained the time scales for *d*(28 − 71), *d*(18 − 203), *d*(122 − 159) and *d*(36 − 129). The values are ≈ 0.3*ms*, ≈ 3.9*ms*, ≈ 0.4*ms* and ≈ 1.7*ms*. The corresponding rates of formation for the NMP domain, global molecule, LID and interface between NMP and LID domains are 3, 472*s*^−1^, 255*s*^−1^, 2, 342*s*^−1^ and 584*s*^−1^. These calculations show that the NMP and LID domains form early in the folding process and their interface forms before collapse. The distributions of these four distances at different times, shown in Fig.S5., provide a more detailed picture of the assembly dynamics of different regions of ADK. The results in Fig.S5 show that there is a great deal of dispersion in the ordering of various parts of the ADK structure.

### Parallel pathways and kinetic intermediates

We calculated the fraction of native contacts of each domain, *Q*^*NMP*^, *Q*^*LID*^, *Q*^*CORE*^, from the 100 folding trajectories. Using these as progress variables for the folding reaction, we find that ADK folds by multiple parallel pathways. The NMP and the LID domains fold cooperatively in a two-state manner, albeit at different rates, while the CORE domain folds through 5 successive stages, which is illustrated using a sample folding trajectory at the bottom right of Fig.5. In each of these stages, the CORE domain becomes increasingly ordered with acquisition of the native-like structure occurring in the final stage. Therefore, ADK could fold through 2 × 2 × 5 =20 states. However, in the 100 folding trajectories, only 13 states are kinetically populated. We classify these as LL1, HL1, LH1, HH1; LL2, HL2, LH2, HH2; HL3, LH3, HH3; HH4; HH5. The first letter in the 3-letter notation represents the state of NMP domain; “L” means the value of *Q*^*NMP*^ is Low and NMP domain is unfolded, “H” means the value of *Q*^*NMP*^ is High and NMP domain is folded. The second letter represents the state of LID domain, and the third letter stands for the state of the CORE domain. Labels “1 − 5” denote different levels for the values of *Q*^*CORE*^. LL1 is the starting unfolded state and HH5 is the final folded state. There are 15 distinct folding pathways found in the generated folding trajectories (See Table.1 in the SI).

**Figure 5:**
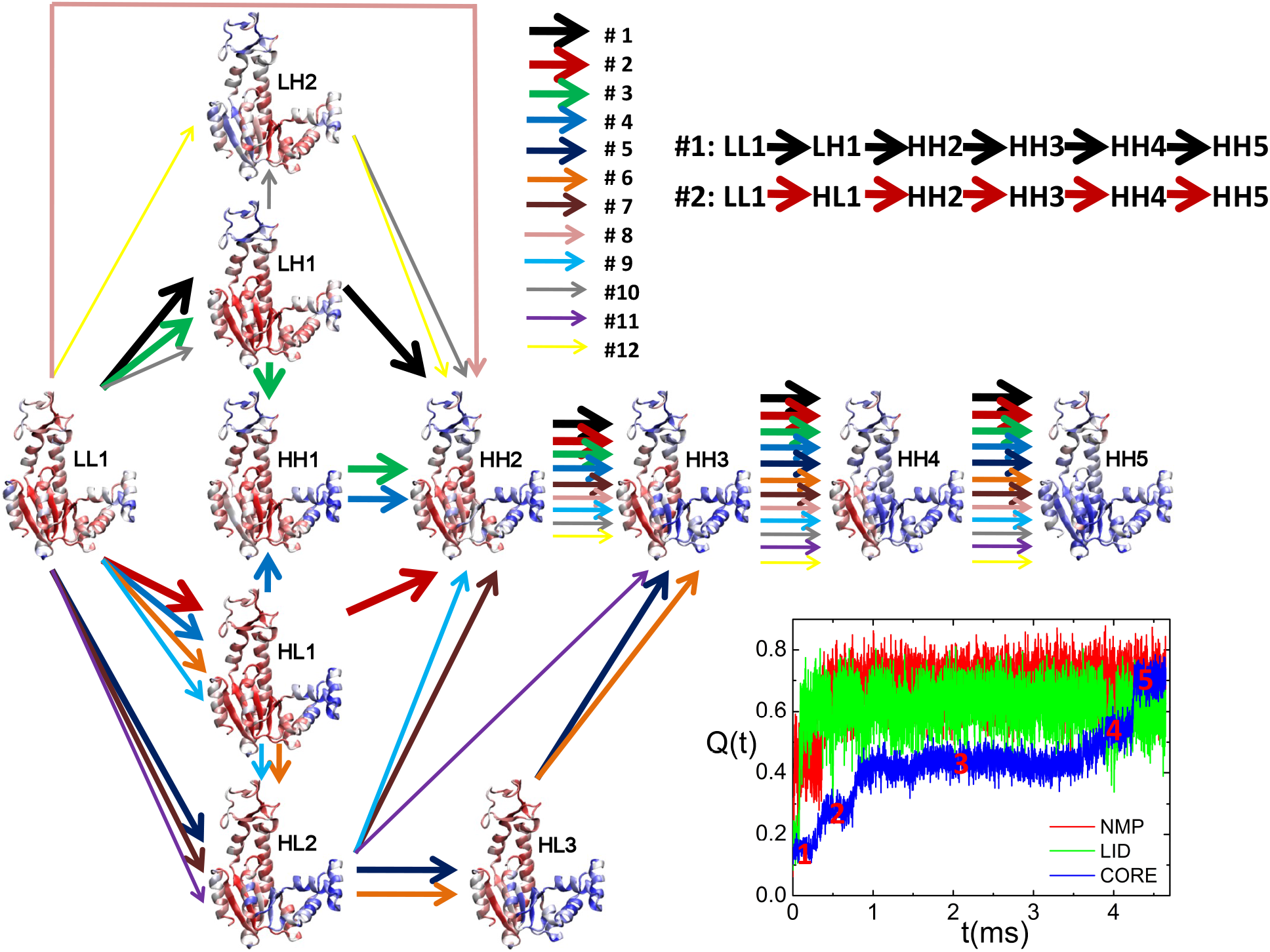
Network of connected states accessed during folding kinetics and parallel pathways. Each state is colored according to the average fraction of the native contacts formed at each residue. Color code: blue, structured; red, unstructured. The 12 most probable folding pathways are represented by the colored arrows with the line widths representing the probability of each folding pathway. The panel on the bottom right shows one representative folding trajectory. The hierarchy of assembly of the domains is clear. The NMP and LID domains form prior to the formation of the CORE domain. A sequence of transitions (1 to 5) drive consolidation of the folded ADK.

The fluxes through the 13 states follow a complex pattern, as illustrated in Fig.5, where each state is colored according to the average fraction of native contacts formed at each residue. As observed in the single-molecule experiment,^5^ and in a previous coarse-grained thermally-triggered folding simulation,^16^ the folding trajectories might involve transitions between the distinct states, thus introducing loops in the folding pathways. For simplicity, we removed these loops from the figure and only the 12 most probable pathways are shown in Fig.5. From this folding flux diagram, we find that the early stage in the folding reaction for ADK is very plastic while the late stage is more restricted, which reflects the narrowing of the folding free energy landscape to the native state. In addition, there is a pathway that directly connects the globally unfolded state (LL1) to HH2 from which folding to the native state (HH5) occurs sequentially (Fig.5). It is likely that the fluxes through the metastable states could be altered by changing the external conditions.^5^

### Thermal and kinetic networks are similar

To illustrate the structural similarities between the 7 thermal substates with significant populations and the 13 substates identified from the kinetic folding trajectories, we computed the average fraction of native contacts formed by every residue, *f*_*Q*_, for the 20 states. For each thermal substate, we searched for the kinetic state that has a high degree of correlation (exceeding 0.9) between thermal and kinetic *f*_*Q*_. For the thermal substates S2 and S9, we could not find suitable matching kinetic states, which means these two substates are not sampled in the kinetic folding trajectories. The other 5 thermal states correlate with the kinetic states (Fig.6). The correlation between S8 and HL3 is very high (R=0.99, Fig.6D). We surmise that S8 and HL3 are structurally similar (in short, *S*8 ∼ *HL*3). Likewise we find *S*11 ∼ *HH*3(R=0.93, Fig.6E), *S*1 ∼ *LL*1(R=0.96, Fig.6A), *S*7 ∼ *HL*1(R=0.98, Fig.6B), *S*12 ∼ *HH*5(R=0.94, Fig.6C). The high degree of correlation for the major states during folding shows that both in thermal and kinetic folding very similar states are sampled. The connectivity between these states, which would define the folding pathways, could vary and may readily be altered by changing the [GdmCl].^5^

**Figure 6:**
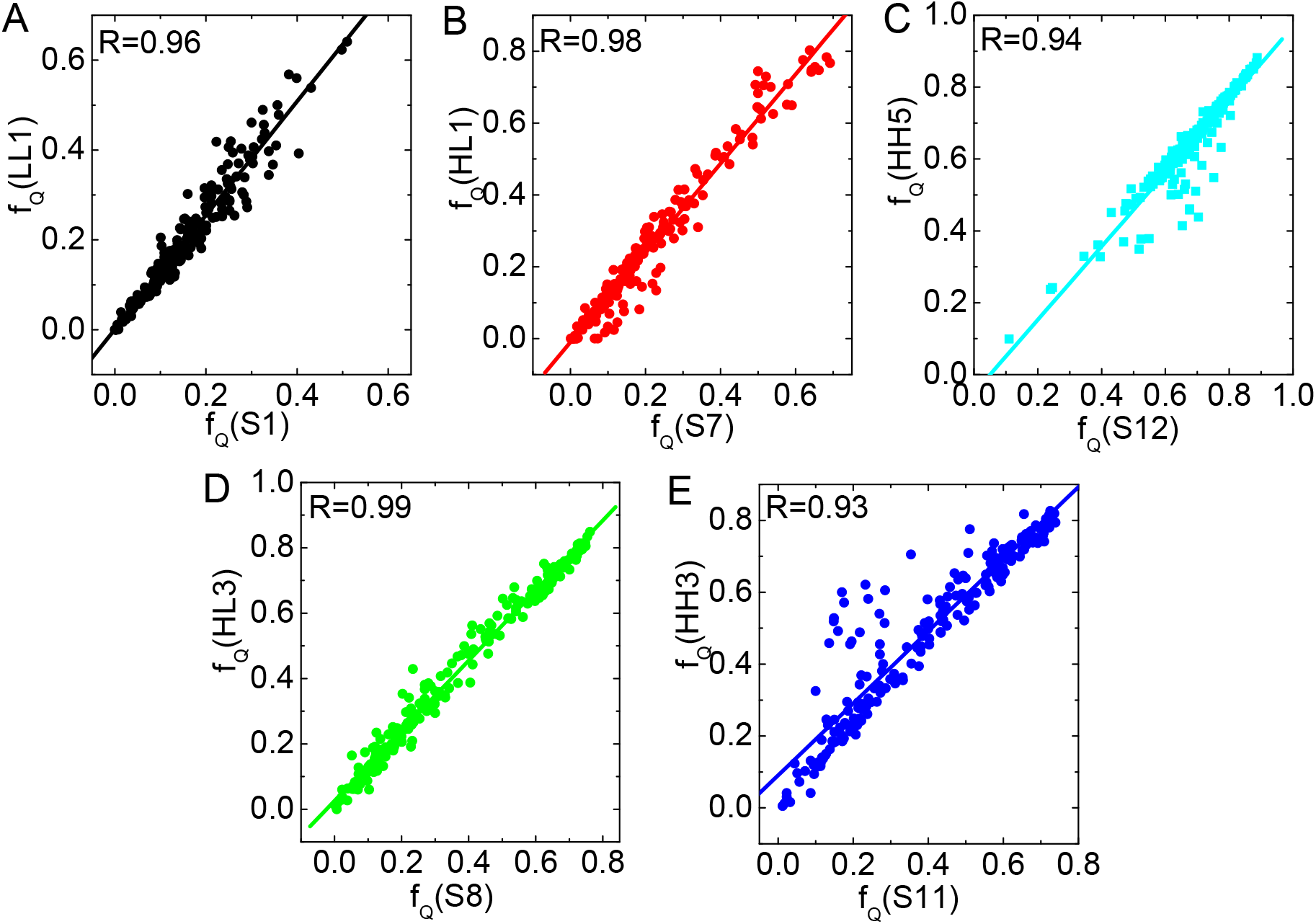
Comparison between thermal and kinetic states. (A) Correlation between *f*_*Q*_s for *S*1 and *LL*1. The correlation coefficient R=0.96. (B) Correlation between *f*_*Q*_s for *S*7 and *HL*1 with R=0.98. (C) Correlation between *f*_*Q*_s for *S*12 and *HH*5 with R=0.94. (D) Correlation between *f*_*Q*_s for *S*8 and *HL*3 with R=0.99. (E) Correlation between *f*_*Q*_s for *S*11 and *HH*3 with R=0.93.

## Discussion

We have shown that the strategy developed here that combines coarse grained SOP-SC model for the multi-domain protein and a phenomenological theory that takes the effects of denaturants into account (GdmCl in the present work) accounts quantitatively for many aspects of folding of ADK, which is an example of MMP. The key results are: (i) The calculated folding times, corrected for stability at the small value of [GdmCl](=0.3M), are in excellent agreement with experiment. (ii) The order of events, as assessed by the time dependent changes in the distances between specific residues (Fig.4C), reproduces the ensemble averaged FRET experiments.^6^ In particular, the finding that the intramolecular distance between residues 28 and 71 reaches the value in the native state extremely rapidly before any global structure acquisition accords well with experiments.^6^ In addition, we find that the slow steps in the consolidation of the native fold involve reduction in the distances between 36-129 (residues in the NMP and LID domains) and between 18-203 reaching the values in the folded state. Although both Q18 and A203 are in the core domain, the rate of approaching the Q18-A203 distance in the folded state is slow, and occurs only upon global folding, a conclusion that is also in accord with ensemble FRET experiments.

### Cooperativity

Communication between domains during the assembly of ADK, as the temperature is decreased, is dramatically different from the all-or-none behavior normally observed in ensemble experiments in single domain proteins^30–35^ or in the folding of SMPs. We expect, based on the structure of ADK (Fig.1C), that the NMP and (possibly) the LID domains could order nearly independently in a two-state manner because their sequences are contiguous in sequence and in the arrangement of the secondary structural elements. This expectation is borne out in the plots in Fig.2A, which shows that the more stable NMP domain melts at the higher temperature 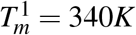. Rao and Gosavi^17^ also came to similar conclusions.

Similarly, a two-state like, albeit with less cooperativity, is observed in the folding of the less stable LID domain whose melting temperature is 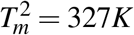. The near independence of their folding is also reflected in the free energy profiles shown in Fig.S3 in the SI. Comparison of Fig.S3A and Fig.S3D shows that the NMP domain forms independently of the LID domain. At both 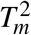 and 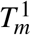 the average value of *Q*^*NMP*^ exceeds 0.5 while the value of *Q*^*LID*^ remains small until the temperature is reduced below 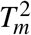 (Fig.2A). In other words, the formation of the NMP domain does not induce order in the LID domain. Similar conclusions may be drawn by comparing the temperature-dependent profiles for *Q*^*NMP*^ and *Q*^*CORE*^ shown in Fig.2A and the free energy profiles shown in Fig.S3B and Fig.S3E. The NMP and CORE domains assemble independently, which indicates that there is little communication between the two domains.

In contrast, both from the perspective of thermodynamics and kinetics, the folding of the LID and CORE domains are intertwined. As the temperature decreases below 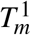, the *Q* for the CORE increases and only when it reaches ≈ 0.5 there is a sharp increase in ⟨*Q*⟩ for the LID (see the green curve in Fig.2A). The results in Fig.S3C and Fig.S3F also suggest that at 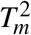 order in the CORE and the LID domain are coordinated in the sense that decrease in the free energy associated with the CORE domain also results in an increase in the stability of the LID domain. In other words, folding of the LID domain is slave to acquisition of certain order in the CORE domain. The importance of cooperative interactions between the LID and CORE domain, which is a consequence of contacts between the reentrant secondary structural elements in the folded state (Fig. S1), was previously established^17^ by comparing the equilibrium energy profiles of ADK and circular permutants. Finally, we note that, even after folding is complete, the LID domain is less stable compared to the other two domains.^23^ The decreased stability of the LID domain might be a conduit to facilitate allosteric transitions.^36,37^

Based on optical tweezer experiments on T4 lysozyme,^4^ which also harbors a reentrant helix like ADK, it has been argued that cooperativity between the distant parts of the chain is needed for stability. It is tempting to speculate that interwoven discontinuous chain topology in MMPs might be an evolutionary consequence not only for stability but also for functional purposes.

### Pathways

Both equilibrium, thermal melting and denaturant-induced unfolding show that multiple states are sampled as the transition from the folded state to the unfolded state occurs in ADK. In addition, the refolding kinetics reveals that an intricately connected network of metastable states is involved in the route to the folded state. The fluxes through these states are dramatically different, which suggests that refolding is heterogeneous. The heterogeneity in the folding pathway has been shown in smFRET experiments,^5,10^ which established that the complexity of the pathways increases as GdmCl concentration increases. The kinetic simulations further support the conclusion reached in experiments. Because smFRET uses only a one-dimensional coordinate for the structures, the metastable structures could not be determined. Our simulations (Fig.5) reveal that the folding pathways are highly complex and heterogeneous, and gives possibly an even more nuanced picture than suggested based on analyses of the experimental data.^5,10^ We find that at the late stages of folding (HH3 → HH4 → HH5), the contacts in the NMP and the LID domain are fully formed but various secondary structural elements in the CORE domain have not fully folded (see the bottom right panel in Fig.5). In the HH3 → HH4 transition the helix given in orange in Fig.1C becomes ordered and in the HH4 → HH5 transition helix displayed in red in Fig.1C gets acquired, resulting in an increase in *Q*^*CORE*^ (see the blue line in the sample folding trajectory in Fig.5).

Although the late stages of folding occur sequentially (HH3→HH4→HH5), folding pathways are heterogeneous before HH3 forms. Fig.5 vividly illustrates that there are multiple routes to the formation of HH3. In some of the pathways the LID domain forms first but in others the NMP domain forms before the LID domain. The network is multiply connected in the sense HH2 can be accessed through LL1→HH2 or by the pathway LL1→LH2→HH2. Such complex network of pathways through which the fluxes could dramatically change does not exist in single domain proteins or possibly in SMPs, Simply connected Multidomain Proteins.

### Rules for the folding of multi-domain proteins

Some general lessons about multi-domain proteins emerge from the current work when integrated with previous studies. (1) The domains in the SMPs assemble almost independently with the stability being determined by interface between neighboring constructs. For instance, in the ankyrin repeat proteins (Fig.1A) folding is triggered by interaction between domains *i* and *i* + 1, which propagates till assembly is complete.^38^ (2) In other homo-oligomeric SMP complexes, the interactions at the interfaces contribute most to the stability, which implies that the nature of residues at the junction of domains must play a key role. From a kinetic perspective, the orientations of the domains are significant, as shown in the assembly of the allosteric tetrameric protein, L-lactate dehydrogenase.^39^ In these cases, the free energy of stability is approximately the sum of the individual domains and an additional gain in the interface formation with correct relative orientation. (3) In contrast, in the MMPs in which the domains are discontinuous from the sequence and SSE perspective, communication between domains is most relevant. That this is the case has been shown in pulling experiments on T4 Lysozyme^4^ in which a reentrant helix stabilizes interaction between the two domains. Similarly, the C-terminal helix *α*9 and the N-terminal helix *α*1 and the strand *β* 3 play the analogous role in ADK (Fig.S1 in the SI). We speculate that the enhanced stability due to the interwoven contacts might be needed to minimize fluctuations in the *apo* folded state (see Supplementary Fig.1 in the pulling experiment^23^) in order to facilitate the closed to open transition in the LID domain, which is required for function. (4) During the refolding kinetics, the continuous domains fold before the discontinuous domains. We do not find that the CORE domain is formed before the other two. The same conclusion was reached in the refolding of the two Dihydrofolate reductase^40^ in which the continuous adenosine loop domain always folds before the discontinuous loop domain. Despite such a stringent requirement in the order of folding of the domains, there are many metastable states that are visited during the early stages of folding (Fig.5), which attests to the plasticity of the folding landscape.^5^

## Concluding Remarks

The general conclusion that emerges from this study is that cooperative interactions in multidomain proteins, with discontinuity in sequence and interactions between reentrant secondary structural elements that stabilize the native fold, arise late in the folding process. The early stages of folding and assembly are highly dynamic. It is likely that similar rules also hold in the self-assembly of ion-driven folding of ribozymes (RNA molecules that function as enzymes), which are composed of many domains.^41,42^ In *Azoarcus* ribozyme several of the domains could fold independently. However, some parts of the sequences are interwoven and cooperative interactions between domains that harbor these domains typically occur only above the midpoint of the ion concentration (usually Mg^2+^). It would be interesting to use single molecule pulling experiments^4^ to dissect inter-domain interactions in RNA.

## Methods

### SOP-SC model

We carried out simulations using the SOP-SC (Self-Organized Polymer-Side Chain) model for the protein.^14,43^ Each residue is represented by two interaction beads with one located at the C_*α*_ position and the other at the center of mass of the side chain. The SOP-SC energy function is

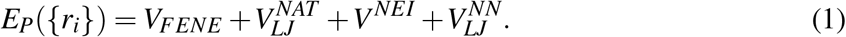

The detailed functional forms for 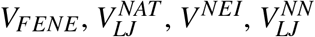 and the values of the parameters are described elsewhere.^14^

### Molecular Transfer Model (MTM)

Currently the MTM is the only available computational method that accurately predicts the outcomes of experiments and provides the structural basis for folding as a function of denaturants^15,44,45^ and pH.^46^ In the MTM, whose theoretical basis is provided in a previous study,^43^ the effective free energy function for a protein in aqueous denaturant solution is given by,

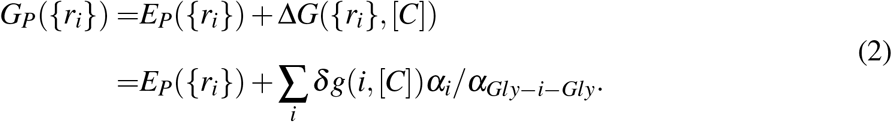

In Eq.(2), Δ*G*({*r*_*i*_}, [*C*]) is the free energy of transferring a given protein conformation from water to an aqueous denaturant solution with [*C*] being the concentration. The sum in the above equation is over all the beads, *δg*(*i*, [*C*]) is the transfer free energy of the interaction center *i, α*_*i*_ is the solvent accessible surface area (SASA) of the interaction center *i*, and *α*_*Gly*−*i*−*Gly*_ is the SASA of the interaction center *i* in the tripeptide *Gly* − *i* − *Gly*. We used the procedure described previously^14,43^ to calculate the thermodynamic properties of proteins in the presence of denaturants.

### Langevin and Brownian Dynamics Simulations

We assume that the dynamics of the protein is governed by the Langevin equation,

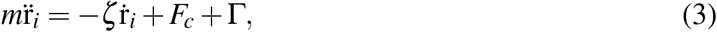

where *m* is the mass of a bead, *ζ* is the friction coefficient, *F*_*c*_ = −*∂E*_*P*_({*r*_*i*_})*/∂r*_*i*_ is the conformational force calculated using Eq. (1), Γ is the random force with a white noise spectrum.

We performed Langevin simulations using a low friction coefficient *ζ* = 0.05*m/τ*_*L*_.^22^ The equations of motions were integrated using the Verlet leap-frog algorithm. To enhance conformational sampling, we used Replica-Exchange Molecular Dynamics (REMD).^19–21^

In order to simulate the folding kinetics, we set *ζ* = 50*m/τ*_*L*_, which approximately corresponds to the value in water.^25^ At the high *ζ* value, we use the Brownian dynamics algorithm^47^ to integrate equations of motion using

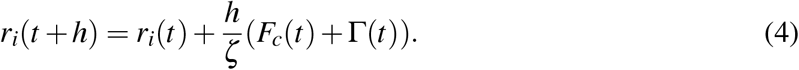

### Data Analysis

We identify the melting temperature as the peak position in the specific heat as a function of temperature, 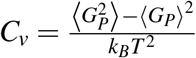.

The structural overlap function, 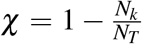,^9^ is employed to monitor the folding/unfolding reaction, where

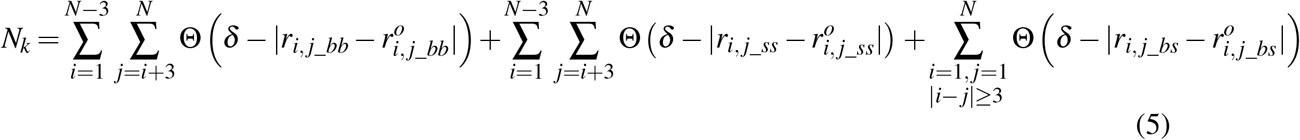

In Eq. (5), Θ(*x*) is the Heavyside function. If 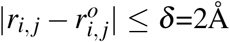, there is a contact. *N*_*k*_ is the number of contacts in the *k*^*th*^ conformation, and *N*_*T*_ is the total number of contacts in the folded state. The microscopic order parameter of the protein, *χ*, is used to distinguish between the native, unfolded and intermediate states.

## Acknowledgement

We are grateful to Prof. Gilad Haran and Dr. David Scheerer for useful comments. ZL acknowledges financial support from the National Natural Science Foundation of China (11104015, 11735005) and the China Scholarship Council (201806045049). DT is grateful to the National Science Foundation (CHE 19-000033), and the Collie-Welch Regents chair (F0019) for supporting this work.

