## supporting information for "Cooperativity and folding kinetics in a multi-domain protein with interwoven chain topology"

Zhenxing Liu

*Department of Physics, Beijing Normal University, Beijing 100875, China**

D. Thirumalai

*Department of Chemistry, The University of Texas at Austin, Austin, TX 78712*†

*Electronic address:

†Electronic address:

**Folding kinetics:** To generate folding trajectories at $[GdmCl] = 0M$ using Brownian dynamics simulations[1], we first prepared 100 trajectories starting **ZL Do you mean starting?**from the native structure at $T = 450K$. These structures were equilibrated them for $10^8$ time steps before quenching the temperature to $T = 293K$ in order to initiate folding. The distribution of first passage times, $P_{fp}(s)$, is calculated using,

$$P_{fp}(s) = \frac{1}{M}\sum_{i=1}^{M}\delta(s - \tau_{1i}) \tag{1}$$

where $M$=100 is the total number of trajectories, $\tau_{1i}$ is the time required for the $i^{th}$ trajectory to reach the Native Basin of Attraction for the first time. The fraction of unfolded molecules at time $t$ is computed using,

$$P_u(t) = 1 - \int_0^t P_{fp}(s)ds. \tag{2}$$

By fitting $P_u(t) = e^{-tk_f}$ the folding rate at $[GdmCl] = 0M$ is extracted.

**Stability-corrected folding rate calculation:** To estimate the folding rate in GdmCl solution, $k_f([GdmCl])$, using the folding rate at [GdmCl]=0, $k_f(0)$, we exploited the following equation,

$$k_f([GdmCl]) = k_f(0)e^{\frac{\Delta G(0)-\Delta G([GdmCl])}{k_BT}}, \tag{3}$$

where $\Delta G$ is the stability of native states with respect to unfolded states. Using the above equation and $\frac{(\Delta G(0)-\Delta G([GdmCl])}{k_BT}$ shown in Fig.S4A, we calculated the dependence of $k_f([GdmCl])$ on $[GdmCl]$ (see Fig.S4B). The red circle is the experimental value[2, 3]. Although the $\Delta G([GdmCl]$ calculated from simulations at $T = 293K$ are larger than the experimental values, the stability-corrected rates ($k_f([GdmCl])$) are in the reasonable range.

[1] Veitshans, T.; Klimov, D.; Thirumalai, D. *Folding and Design* **1997**, *2*, 1–22.

[2] Ishay, E.; Rahamim, G.; Orevi, T.; Hazan, G.; Amir, D.; Haas, E. *J. Mol. Biol.* **2012**, *423*, 613–623.

[3] Ratner, V.; Amir, D.; Kahana, E.; Haas, E. *J. Mol. Biol.* **2005**, *352*, 683–699.

Table 1: 15 distinct folding pathways and the associated flux, expressed as probability in %.

| Pathway# | Pathway | Probability % |
|---|---|---|
| 1 | LL1→ LH1→ HH2→ HH3→ HH4→ HH5 | 14 |
| 2 | LL1→ HL1→ HH2→ HH3→ HH4→ HH5 | 14 |
| 3 | LL1→ LH1→ HH1→ HH2→ HH3→ HH4→ HH5 | 12 |
| 4 | LL1→ HL1→ HH1→ HH2→ HH3→ HH4→ HH5 | 10 |
| 5 | LL1→ HL2→ HL3→ HH3→ HH4→ HH5 | 10 |
| 6 | LL1→ HL1→ HL2→ HL3→ HH3→ HH4→ HH5 | 7 |
| 7 | LL1→ HL2→ HH2→ HH3→ HH4→ HH5 | 7 |
| 8 | LL1→ HH2→ HH3→ HH4→ HH5 | 5 |
| 9 | LL1→ HL1→ HL2→ HH2→ HH3→ HH4→ HH5 | 5 |
| 10 | LL1→ LH1→ LH2→ HH2→ HH3→ HH4→ HH5 | 4 |
| 11 | LL1→ HL2→ HH3→ HH4→ HH5 | 4 |
| 12 | LL1→ LH2→ HH2→ HH3→ HH4→ HH5 | 3 |
| 13 | LL1→ LL2→ LH2→ LH3→ HH3→ HH4→ HH5 | 2 |
| 14 | LL1→ LL2→ LH2→ HL3→ HH3→ HH4→ HH5 | 2 |
| 15 | LL1→ LL2→ HL3→ LH3→ HH3→ HH4→ HH5 | 1 |

Table 2: Fraction of native contacts ($f_Q$) formed in the 13 kinetic states. HH5 is the folded state. The meaning of the states, such as HL3, is explained in the main text.

| State# | State Name | $f_Q$ |
|---|---|---|
| 1 | LL1 | 0.17 |
| 2 | HL1 | 0.23 |
| 3 | LH1 | 0.24 |
| 4 | HH1 | 0.31 |
| 5 | LL2 | 0.26 |
| 6 | HL2 | 0.33 |
| 7 | LH2 | 0.35 |
| 8 | HH2 | 0.40 |
| 9 | HL3 | 0.42 |
| 10 | LH3 | 0.43 |
| 11 | HH3 | 0.50 |
| 12 | HH4 | 0.57 |
| 13 | HH5 | 0.68 |

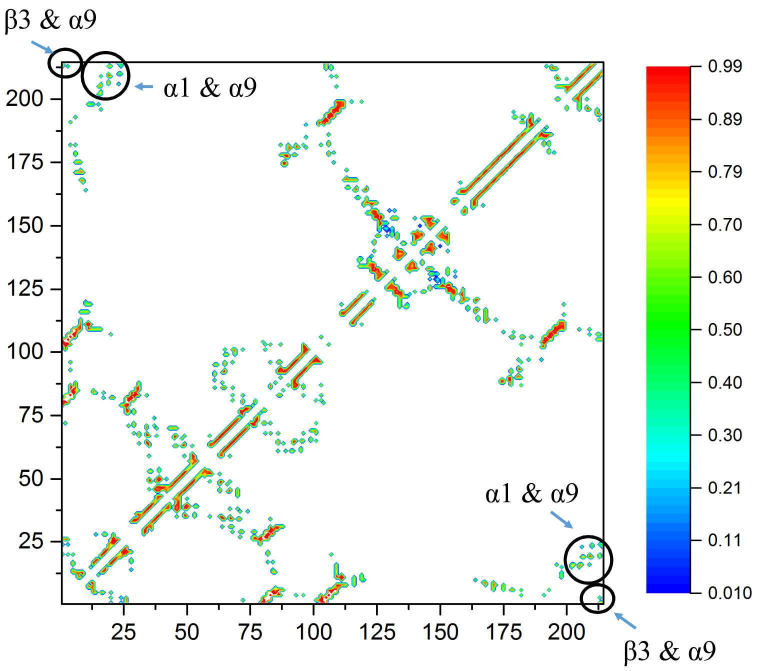


FIG. S1: Contact map of the native state ensemble at 293K for ADK. The reentrant C-terminal helix, $\alpha 9$, forms contacts with the N-terminal helix $\alpha 1$ and strand $\beta 3$ (shown in circles). The residues spanning $\alpha_9$, $\alpha_1$, and $\beta_3$ are 202-213, 13-24, 2-7.

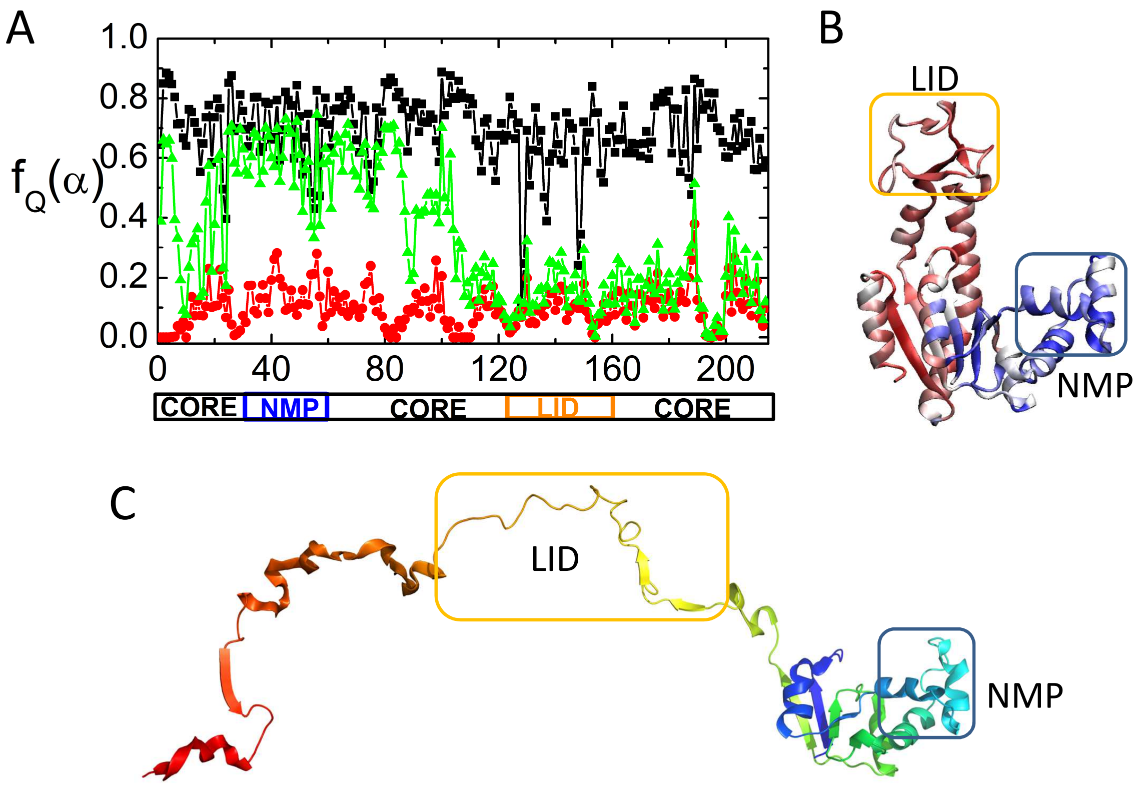


FIG. S2: Structural features of the intermediate state $I^G$. (A) The average fraction of the native contacts formed at each residue for the global states, $f_Q(\alpha)$, where $\alpha$ labels the native state $N^G$ (black line), unfolded state $U^G$ (red line), and intermediate state $I^G$ (green line), respectively. (B) $I^G$ is colored according to the average fraction of the native contacts formed at each residue. Color code: blue, structured; red, unstructured. (C) A representative structure for $I^G$ is shown.

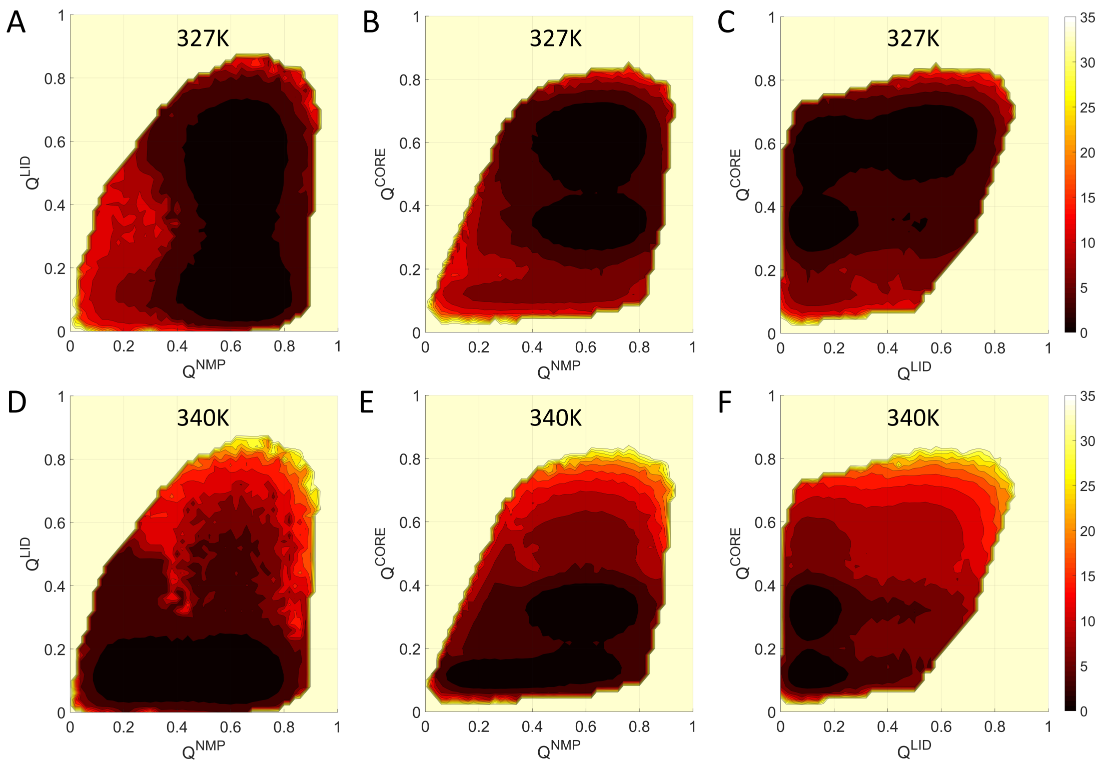


FIG. S3: Two dimensional free energy profiles. We subtracted the minimum value of the free energy from each panel. The scale on the right (in kcal $\cdot$ mol$^{-1}$) indicates that smaller values are closer to the minimum. (A) $G(Q^{NMP}, Q^{LID})$ at $T_m^2$. (B) $G(Q^{NMP}, Q^{CORE})$ at $T_m^2$. (C) $G(Q^{LID}, Q^{CORE})$ at $T_m^2$. (D) $G(Q^{NMP}, Q^{LID})$ at $T_m^1$. (E) $G(Q^{NMP}, Q^{CORE})$ at $T_m^1$. (F) $G(Q^{LID}, Q^{CORE})$ at $T_m^1$. The values of the melting temperatures are $T_m^2 = 327K$ and $T_m^1 = 340K$.

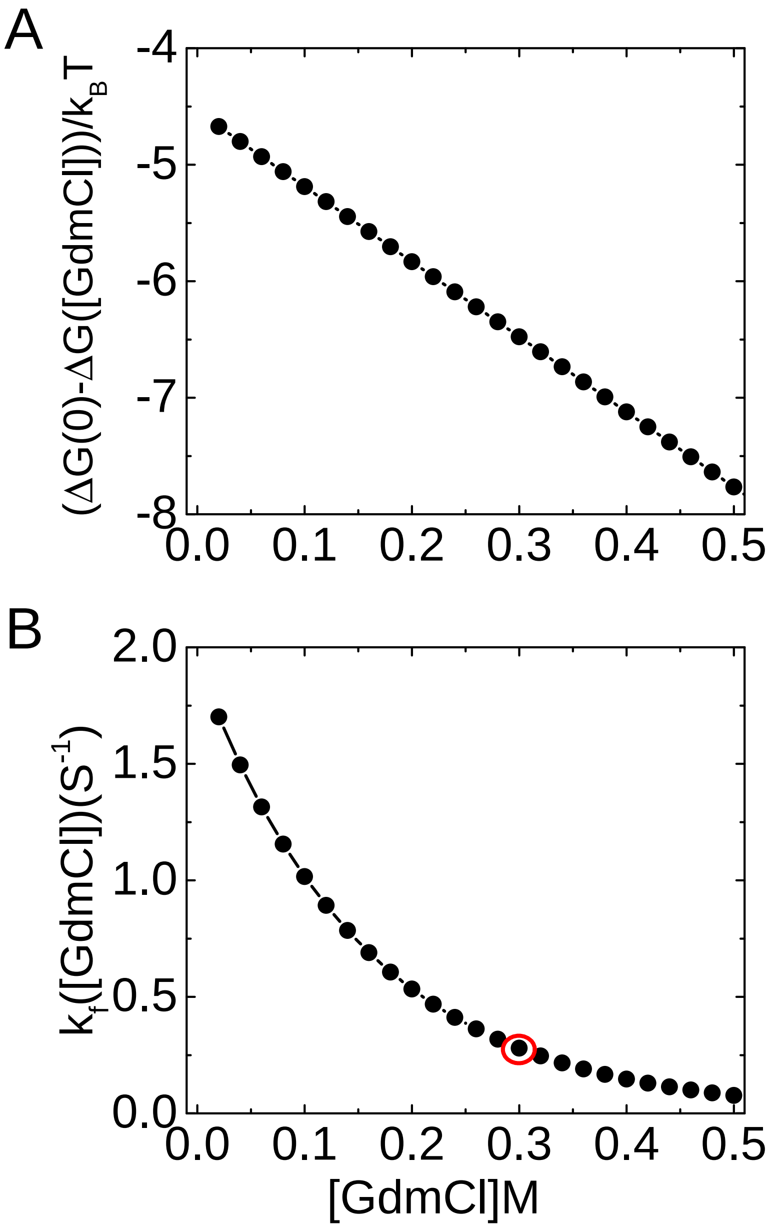


FIG. S4: (A) Relative changes in the free energy of stability of the folded ADK with respect to the unfolded state as a function of [GdmCl]. (B) The estimated folding rate (Eq.(3)) as a function of [GdmCl], $k_f([GdmCl])$. The red circle at 0.3M is the value of $k_f([0.3M])$ measured in experiments [2].

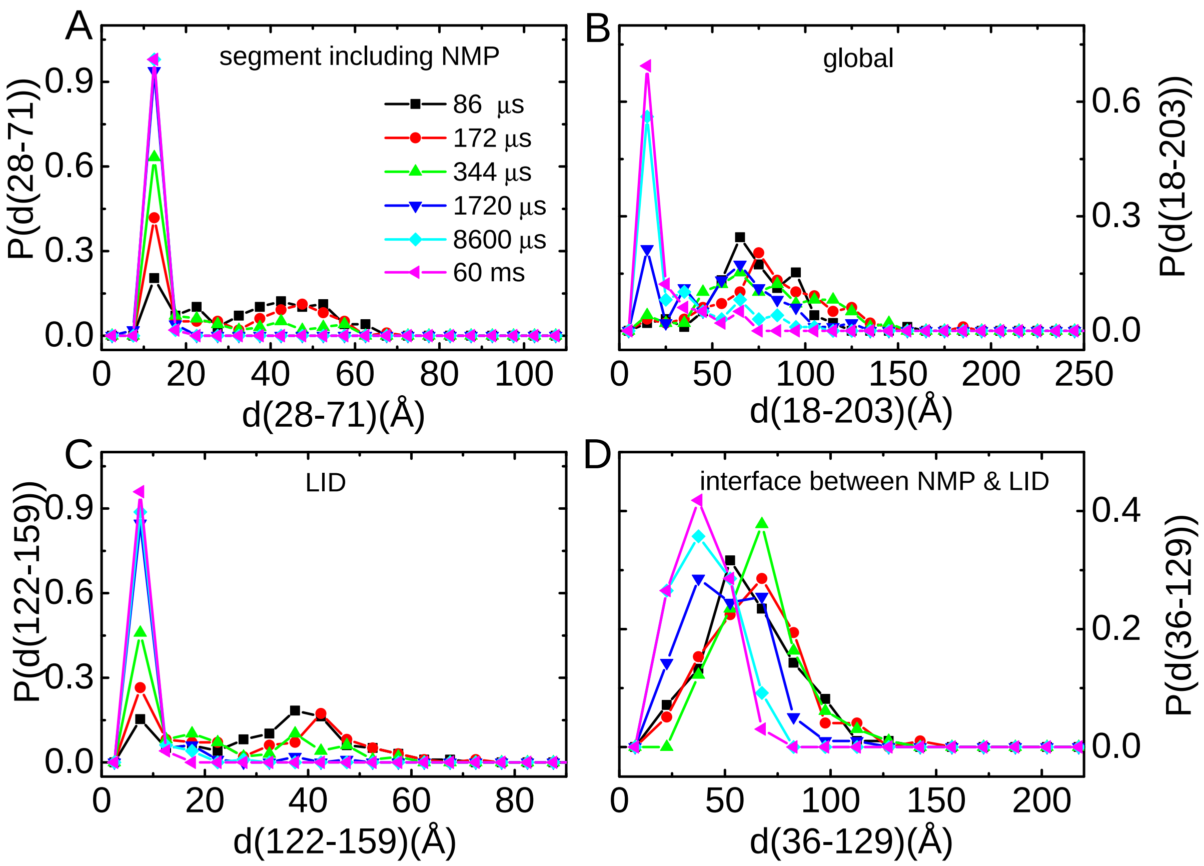


FIG. S5: Distributions of distances between four labeled pairs of residues at times, 86$\mu s$, 172$\mu s$, 344$\mu s$, 1720$\mu s$, 8600$\mu s$ and 60$ms$ along the kinetic folding trajectories. (A) Residue pair 28 and 71. (B) Residue pair 18 and 203. (C) Residue pair 122 and 159. (D) Residue pair 36 and 129. In the folded state, the distances between these four pairs of residues are 11.3Å, 13.1Å, 7.6Å, and 26.5Å, respectively.
